# Evaluating species distribution models with discrimination accuracy is uninformative for many applications

**DOI:** 10.1101/684399

**Authors:** Dan L. Warren, Nicholas J. Matzke, Teresa L. Iglesias

**Author notes:** Author Contributions: All authors contributed to study design and manuscript preparation. Simulations were coded and run by DLW, literature review was performed by TLI. Post hoc analyses were performed by TLI and DLW.

## Abstract

**Aim:** Species distribution models are used across evolution, ecology, conservation, and epidemiology to make critical decisions and study biological phenomena, often in cases where experimental approaches are intractable. Choices regarding optimal models, methods, and data are typically made based on discrimination accuracy: a model’s ability to predict subsets of species occurrence data that were withheld during model construction. However, empirical applications of these models often involve making biological inferences based on continuous estimates of relative habitat suitability as a function of environmental predictor variables. We term the reliability of these biological inferences “functional accuracy.” We explore the link between discrimination accuracy and functional accuracy.

**Methods:** Using a simulation approach we investigate whether models that make good predictions of species distributions correctly infer the underlying relationship between environmental predictors and the suitability of habitat.

**Results:** We demonstrate that discrimination accuracy is only informative when models are simple and similar in structure to the true niche, or when data partitioning is geographically structured. However, the utility of discrimination accuracy for selecting models with high functional accuracy was low in all cases.

**Main conclusions:** These results suggest that many empirical studies and decisions are based on criteria that are unrelated to models’ usefulness for their intended purpose. We argue that empirical modeling studies need to place significantly more emphasis on biological insight into the plausibility of models, and that the current approach of maximizing discrimination accuracy at the expense of other considerations is detrimental to both the empirical and methodological literature in this active field. Finally, we argue that future development of the field must include an increased emphasis on simulation; methodological studies based on ability to predict withheld occurrence data may be largely uninformative about best practices for applications where interpretation of models relies on estimating ecological processes, and will unduly penalize more biologically informative modeling approaches.

---

Species distribution models (SDM, alternatively environmental niche models or ENM) use data on species occurrences in conjunction with environmental data to generate statistical models of species’ ecological tolerances, environmental limits, and potential to occupy different geographic areas. These methods have been used since the 1920s (Cook 1925, Sutherst 2014), but recent years have seen rapid growth in the number of studies employing SDM in fields including ecology, conservation biology, evolutionary biology, and epidemiology (Peterson, Soberón et al. 2011, Coro, Pagano et al. 2013, Allen and Lendemer 2016, Gutierrez-Tapia and Palma 2016, Lezama Ochoa, Murua et al. 2016, Raghavan, Goodin et al. 2016, Guisan, Thuiller et al. 2017). The primary appeal of SDMs is their tractability; estimating environmental tolerances experimentally is expensive and time-consuming at best, and impractical for many species. In contrast, SDMs can be constructed with minimal investment of resources, using freely available data and software (Hijmans, Cameron et al. 2005, Phillips, Anderson et al. 2006, Thuiller, Lafourcade et al. 2009, Hijmans, Phillips et al. 2012, Kriticos, Webber et al. 2012). For many species of conservation concern, they are one of the only tractable means of estimating habitat suitability, often in cases where stakeholders need these estimates urgently (Keith, Mahony et al. 2014, Warren, Wright et al. 2014).

SDM construction involves many decisions which may affect model predictions. These include choice of modeling algorithm, required sample size, optimal model complexity, choice of study area from which data are drawn, the exclusion of outliers, and selection of environmental predictors, among others (Guisan, Graham et al. 2007, Wisz, Hijmans et al. 2008, Acevedo, Jimenez-Valverde et al. 2012, Domisch, Kuemmerlen et al. 2013, Boria, Olson et al. 2014, Varela, Anderson et al. 2014, Garcia-Callejas and Araujo 2016, Soley-Guardia, Gutierrez et al. 2016, van Proosdij, Sosef et al. 2016). The literature surrounding these decisions is large and growing rapidly, as is the suite of associated software tools. Decisions about how best to model species are typically made using metrics that test discrimination accuracy on subsets of species occurrence data that have been withheld during model construction (Elith, Graham et al. 2006, Radosavljevic and Anderson 2014). However, the binary prediction of withheld occurrence data is rarely the intended application of SDMs; they are more frequently used to make continuous estimates of habitat suitability, and to make predictions outside of the training conditions both in space and in time. These applications often implicitly assume that there is biological meaning to the continuous suitability scores produced by the model, or to the functional relationship between environmental gradients and habitat suitability. However, it is often not clear which (if any) measurable biological phenomena should be correlated with suitability estimates from SDMs. Many of the measurable phenomena that are potentially related to suitability (e.g., population density (Carrascal, Aragon et al. 2015), upper limit of local abundance (VanDerWal, Shoo et al. 2009, Gomes, IJff et al. 2018)) have not been quantified in detail for many real species and as such are unavailable for model validation.

This impracticality of studying environmental suitability experimentally makes it difficult to measure the ability of SDMs to correctly make continuous estimates of habitat suitability. As such, modeling decisions are typically predicated on an assumed relationship between a model’s ability to make continuous estimates of relative habitat suitability (hereafter referred to as “functional accuracy”) and its ability to predict withheld occurrence data (discrimination accuracy). This assumption has been questioned before (Lobo, Jimenez-Valverde et al. 2008), but its importance and validity for SDM studies is largely untested.

Discrimination accuracy is known to be a potentially misleading measure for many applications; it is known to be a poor indicator of model calibration (Reineking and Schröder 2006, Jimenez-Valverde, Acevedo et al. 2013), and may even be negatively correlated with calibration and functional accuracy under some conditions (Murphy and Winkler 1992). This general statistical problem may be exacerbated by attributes of the SDM process in a number of ways. First, spatial autocorrelation present in species distributions and in the environment can generate spurious correlations that a model might treat as biological truth, resulting in models that produce high discrimination accuracy even when occurrence data is random (Raes and ter Steege 2007) or the predictors are biologically meaningless (Bahn and McGill 2007, Bahn and McGill 2013, Fourcade, Besnard et al. 2018). Second, there are phenomena other than the suitability of habitat that shape species distributions (e.g., historical biogeography, dispersal, biotic interactions, Figure 1) (Soberon and Peterson 2005, Kearney 2006, Anderson 2012, Warren 2012, Warren 2013, Warren, Cardillo et al. 2014). Although it is possible to include these processes as predictors for SDMs, this is not often done in practice. Failure to explicitly consider these processes introduces spurious correlations between species occurrences and the environment into the estimate of the environmental niche.

**Figure 1.**
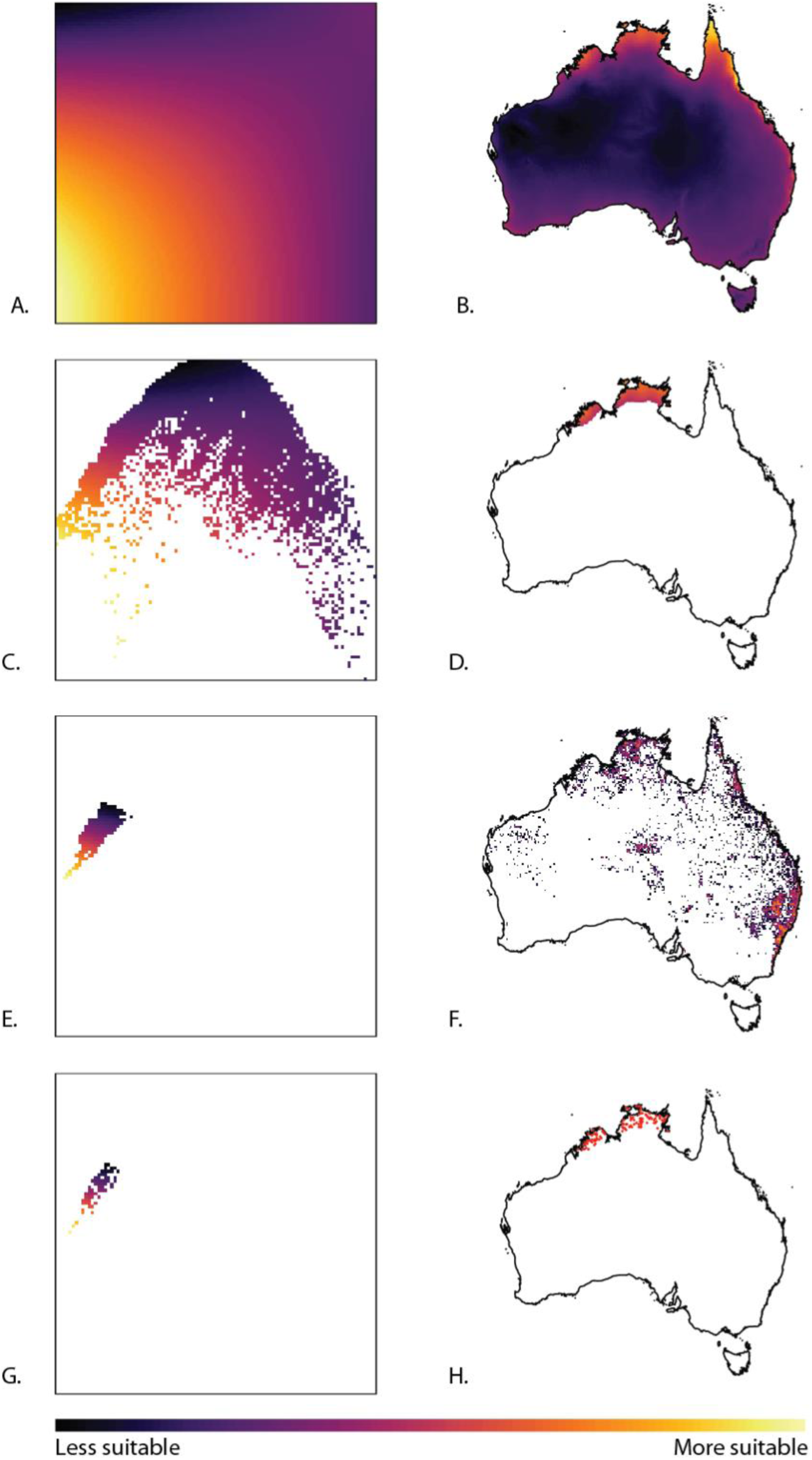
Phenomena affecting species distributions and inference of suitability of habitat. Panel A depicts the niche of a simulated species in the first two principal component axes of the 19 Bioclim variables for Australia. Panel B represents the distribution of suitable habitat for the simulated species. The available habitat present across the continent of Australia only represents a subset of the possible niche space (C). The species’ current range only encompasses a subset of the suitable habitat (D), which further limits the potential distribution of data in environment space (E). Spatial sampling bias (F, see methods) contributes further bias to the representation of the species both in environment space (G) and geographic space (H). While the geographic distribution of the data (H, red points) may resemble the current range of the species (D), the distribution of that data in environment space (G) is a poor representation of the species’ true niche (A). As a result, it may be relatively easy to achieve accurate predictions on randomly withheld occurrence data while still producing a poor estimate of the underlying biology and suitability of habitat.

Similarly, the collection of occurrence data often shows spatial biases (Figure 1, panel F), which may be correlated with spatially autocorrelated predictors (Phillips, Dudik et al. 2009). All of these phenomena can lead to poor niche estimates (Figure 2) that still have high discrimination accuracy in geographic space. Since these non-target phenomena are shared between training and test data, a model that parameterizes the environmental correlates of these processes may have higher discrimination accuracy than a model that accurately estimates the species’ environmental tolerances, and yet may produce pathological behavior in applications where model transferability or continuous estimates of habitat suitability are desired (Lobo, Jimenez-Valverde et al. 2008, Veloz 2009, Radosavljevic and Anderson 2014, Torres, Sutton et al. 2015, Huang and Frimpong 2016).

**Figure 2.**
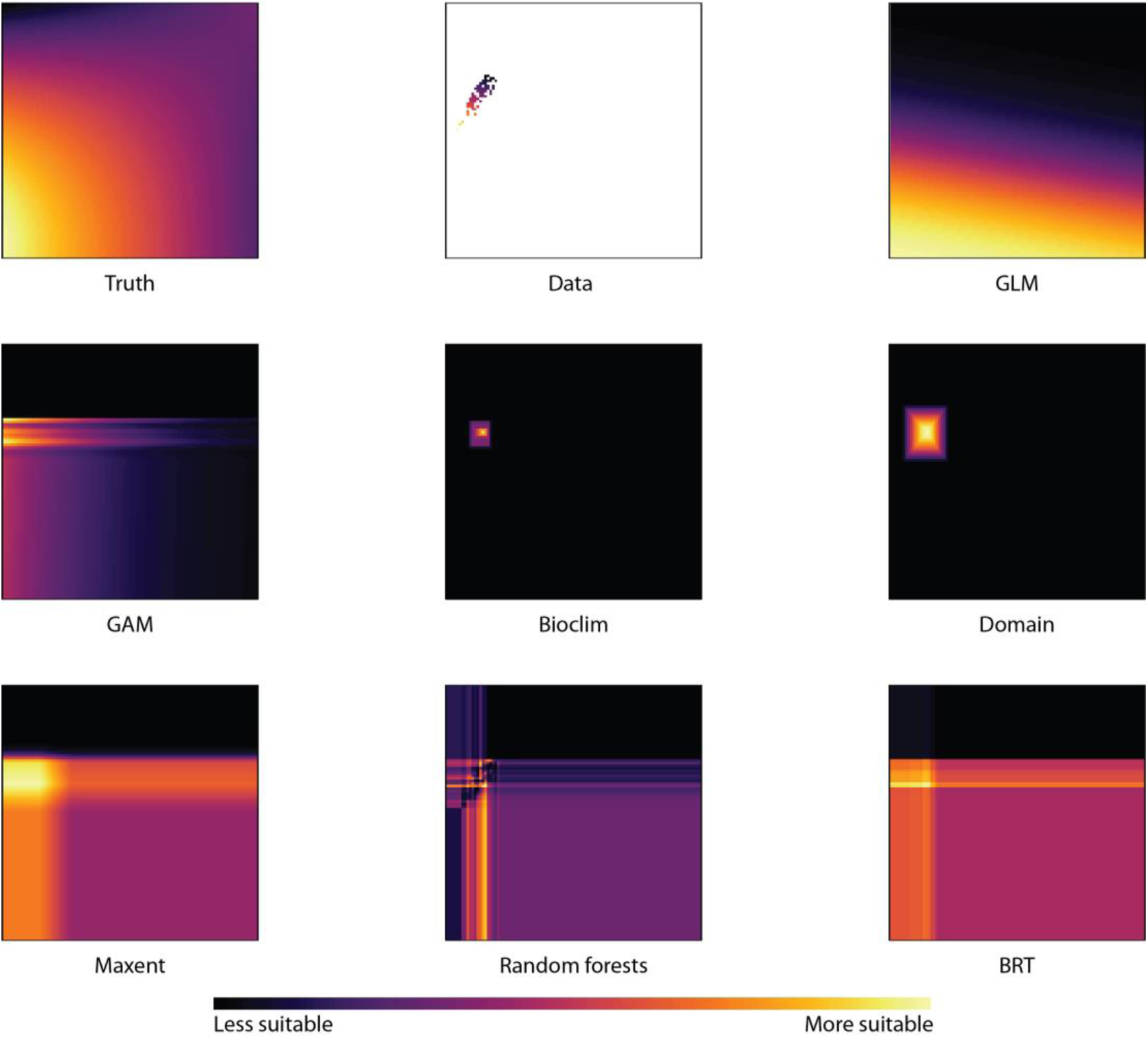
Projection of modeling algorithms in environment space. Using 100 occurrence points for the simulated species in figure 1, we built models using the seven algorithms employed in this study and projected them into the same two dimensional principal component space. The lowest AUC score on 20 randomly withheld data points belonged to random forests (AUC = 0.55), while the highest came from domain (AUC = 0.73). The top left and top center panels show the true niche of the simulated species and the environmental distribution of the data, respectively.

A further issue with discrimination accuracy is the lack of true absence data. One of the primary reasons that SDM methods are so tractable is that they can be used without true absence data, which is often difficult and expensive to obtain. SDMs deal with the lack of true absences by sampling “pseudoabsence” or “background” points which are ideally intended to represent the set of environmental conditions that are potentially accessible to the species. This requires users to make decisions about the size of the appropriate study area for background samples (Acevedo, Jimenez-Valverde et al. 2012), as well as the nature of sampling (e.g., random points or points from closely related species (Phillips, Dudik et al. 2009)). These decisions are often somewhat arbitrary (e.g., background areas chosen using political boundaries or poorly-justified assumptions about dispersal), and can affect both the inferred model (Acevedo, Jimenez-Valverde et al. 2012) and the performance of metrics used to evaluate models (Acevedo, Jimenez-Valverde et al. 2012, Hijmans 2012, Jimenez-Valverde, Acevedo et al. 2013). The lack of real absence data results in models that are incapable of accurately predicting prevalence, and that incorrectly treat some suitable conditions as unsuitable.

Finally, the usefulness of discrimination accuracy as a criterion for selecting SDMs may also be negatively impacted by model complexity. Discrimination accuracy only measures whether a model assigns higher suitability values to presence points than it does to background or absence points, and highly flexible algorithms may produce a broad range of marginal suitability functions that have similar, or even identical, discrimination accuracy (Figure 3). This phenomenon is likely compounded by the frequent use of large numbers of predictors that are highly collinear; as the number of predictors and the complexity of marginal suitability functions increase, the number of potential models with similar discrimination accuracy grows very rapidly.

**Figure 3.**
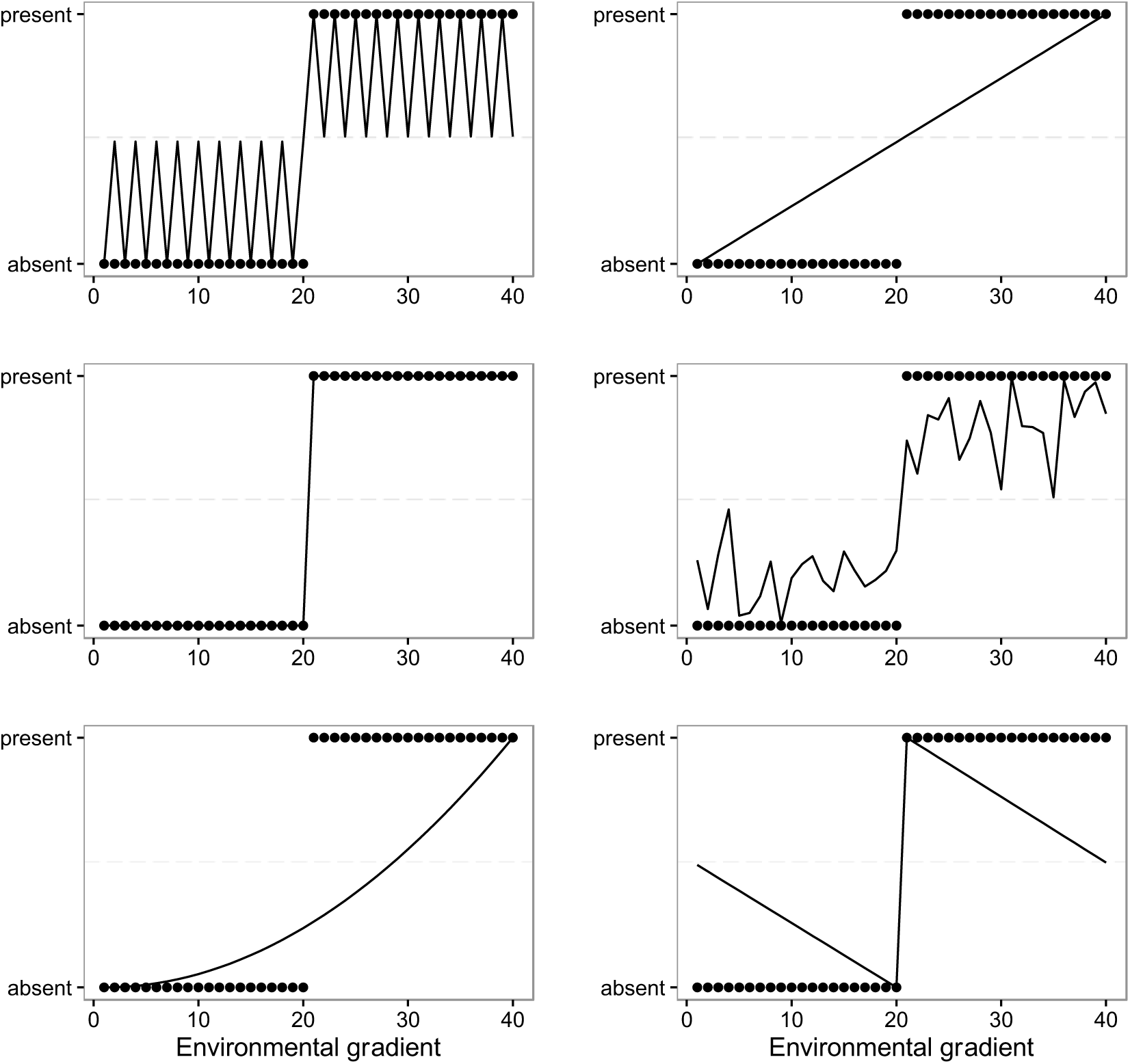
Low information content of discrimination accuracy for inferring functional accuracy. In the above plots, we have simulated 20 absence and 20 presence points along a hypothetical environmental gradient. The six panels represent six hypothetical functions that might be inferred using this data. Each function assigns a higher suitability score (y axis) to all of the presence points than it does to any of the background or absence points. As a result, each function has perfect discrimination accuracy. All six functions are therefore indistinguishable from each other based on discrimination metrics (AUC, TSS, Kappa), while making very different estimates of the functional relationship of habitat suitability to the environmental predictor variable.

Although many of these problems with discrimination accuracy have been noted before, the utility of discrimination metrics for SDM studies has not been examined in a system where the true niche and habitat suitability are known. As a result, we have little information on how useful these metrics are for empirical studies where the goal is to estimate the relative suitability of habitat, despite the ubiquity of discrimination metrics in SDM model selection.

Here we adopt a simulation approach to explore the relationship between discrimination and functional accuracy using virtual species for which the true niche is known. We build models using a number of different algorithms, study area sizes, and methods of partitioning training and test data. However, these simulations are not intended to represent all possible modeling approaches. The goal of these simulations is not to determine which method produces the best niche or distribution estimates, but rather to evaluate commonly used methods for model selection across a broad range of models in a system where we know the underlying true habitat suitability.

## Method

To examine how model selection was affected by modeling algorithm, sampling bias, and non-target spatial phenomena, we conducted four sets of simulation experiments:

1. High complexity: Artificial niches were generated using three randomly chosen variables and models were constructed using all nineteen bioclimatic variables. Background data were drawn from 100 km circular buffers around occurrence points.
2. Geographic partitioning: Same conditions as (1) but presence and background points for each species were split into four quadrants using ENMEval (Muscarella, Galante et al. 2014). Presence and background data from one randomly selected quadrant were withheld for model evaluation.
3. Large background: Same conditions as (1), but background data were drawn from 1000 km circular buffers around occurrence points.
4. Low complexity: Niches were based on two randomly chosen variables and models were built using those two variables plus two more chosen at random.

All analyses used CliMond data for 19 bioclimatic variables typically employed in SDM studies (Nix 1986)_ENREF_1, including data for the present and for 24 combinations of future emissions scenario (A1B and A2), year (2030, 2050, 2070, 2080, 2090, and 2100), and climate model (CSIRO-Mk3.0 and MIROC-H) (Kriticos, Webber et al. 2012). Simulations and analyses were restricted to Australia, including Tasmania.

Simulated niches were created using the generateRandomSp function in the virtualspecies R package (Leroy, Meynard et al. 2015). Simulated species with fewer than 400 suitable grid cells in the initial presence/absence raster showed a strong tendency to produce models that showed no suitable habitat on future climate scenarios, rendering comparisons that were uninformative for model selection. As a result, these simulations were discarded.

To simulate the effects of non-target spatial processes (e.g., historical biogeography, biotic interactions, dispersal limitation), the initial presence/absence raster from virtualspecies was converted to point data, which was partitioned into allopatric regions using k-means clustering. Solutions ranging from 2 to 10 clusters were considered, and an algorithm was used to maximize the minimum distance between clusters. One region was assigned at random to be the range of the species, and converted back into a raster. We recorded the total proportion of suitable cells that fell within this range, to measure the extent to which species distributions departed from the distribution of available suitable habitat across the entire study area.

Spatial sampling bias was modeled using data from 5,969,252 collection records for 28,286 Australian plant species. These records were harvested from GBIF (GBIF.org 2015) using the rgbif (Chamberlain, Boettiger et al. 2013) package, and converted to a raster representing the number of observations per grid cell at the same extent and resolution as the environmental data. All values were then divided by the maximum cell value, resulting in a range of sampling intensity from 0 to 1.

Species occurrence data was sampled from a raster with values in each grid cell *x* calculated as:

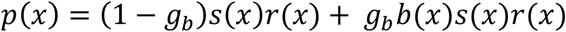

Where probability of sampling is a function of *p(x)*, *gb* is a parameter that controls the magnitude of spatial sampling bias, *b(x)* is the relative strength of spatial sampling bias in cell *x*, *s(x)* is the suitability of habitat in the grid cell, and *r(x)* is a binary variable taking the value 1 inside the species range and 0 everywhere else. For each species, we drew 100 simulated occurrence points by selecting grid cells at random and sampling occurrences as a Bernoulli trial with probability of success equal to *p(x)*. Presence points were drawn with replacement so that we could study the effects of sampling bias. We simulated data across eleven levels of spatial sampling bias, with the bias strength parameter ranging from 0 to 1 in increments of 0.1. We performed 20 simulations for each of the 11 levels of spatial sampling bias. Each of the four simulation conditions (experiments 1-4, above) therefore consisted of 220 simulations, for a total of 880 total simulated species across all experiments.

As noted in a recent review (Meynard, Leroy et al. 2019), simulation studies need to choose both the simulated niches and sampling regimes that are appropriate for the question involved. Since our goal here is to test which metrics select models that accurately estimate the niche, it was essential for us to generate data that would be capable of producing accurate niche estimates in ideal conditions. Due to these concerns we chose not to apply a threshold minimum suitability score below which the organism could not possibly occur; the application of such a threshold would truncate the response functions we are trying to estimate (Meynard, Leroy et al. 2019), resulting in lower expected functional accuracy. Additionally, prior work with virtual species has demonstrated that the application of thresholds results in discrimination accuracy metrics that are overly optimistic (Meynard and Kaplan 2013).

We built models using seven algorithms; Bioclim, Domain, generalized linear models (GLM), generalized additive models (GAM), Maxent, random forests, and boosted regression trees. Models were built using the dismo R package (Hijmans, Phillips et al. 2012) and Maxent (Phillips, Anderson et al. 2006). This resulted in 6160 inferred models, seven for each of the 880 simulated niches. Algorithm settings were left at their default values. For each model, 25 occurrence points were withheld from model construction and used for model evaluation. Each model’s discrimination accuracy was evaluated using three statistics: the area under the receiver operating characteristic curve (AUC) (Fielding and Bell 1997), the true skill statistic (TSS) (Allouche, Tsoar et al. 2006), and Cohen’s kappa (Cohen 1960). While AUC can be calculated using continuous suitability scores, TSS and kappa require binary predictions of presence and absence, so the values for these models that were used in model assessment corresponded to their maximum value across all potential thresholds, i.e., the best possible performance of a thresholded model (Fielding and Bell 1997).

Models were projected onto the current distribution of the environmental variables used for model construction. Models were also projected onto the 24 future climate scenarios. We used the simulated niche from the virtualspecies object to project the true suitability of habitat across those same set of future environments, to assess whether discrimination accuracy is a useful predictor of models’ ability to extrapolate to new environmental conditions. To measure functional accuracy, we compared geographic projections of habitat suitability from the true niche and the inferred models using Spearman rank correlation. Correlations between projected and true suitability scores for the present and for future climate scenarios were measured separately within the species range (areas where *r(x)* = 1) and across the entire study area (Australia and Tasmania). Spearman rank correlation was chosen as a measure of functional accuracy for this study due to the structural differences between models produced by different algorithms, and in consideration of how SDMs are often applied; any two models that assign identical rankings to a set of habitat patches are effectively interchangeable for applications where models are thresholded, or where suitability scores are used to prioritize one habitat patch over another. Rank correlation will reflect this when models produce similar rankings of relative habitat suitability (e.g., ρ = 1 when predictions made from one model are a monotonically increasing function of predictions made from another model). In contrast, Pearson product moment correlation will only assign a value of 1 if the relationship between suitability scores for the two models is linear, which may serve to exaggerate differences that are not relevant to many empirical studies. In order to test the sensitivity of our results to this choice, we also conducted a separate set of analyses using Pearson product moment correlation as a measure of functional accuracy (Appendix S4), but results from these analyses were effectively the same as those seen in tables 1 and 2.

**Table 1.**
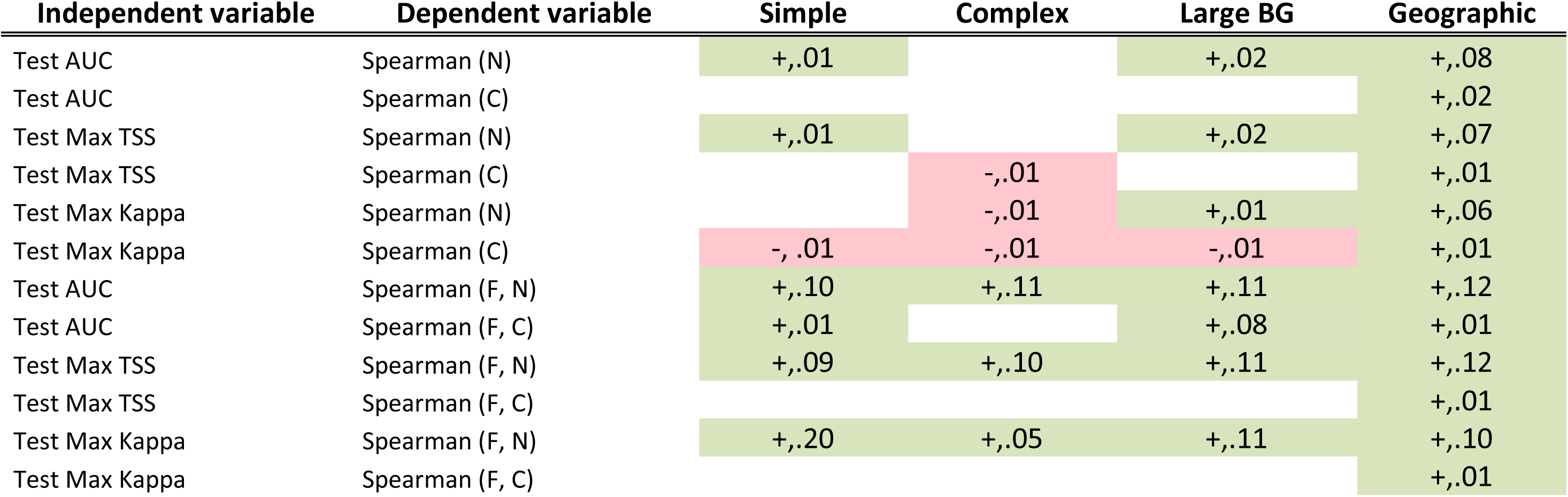
Results of regressions functional accuracy on discrimination accuracy, all algorithms considered together. Significant positive correlations are represented by “+” and green cell color, negative correlations by “-“ and pink cell color. Numbers indicate r^2^ values for each regression. Variables accompanied by (F) indicate that they were measured on models projected across 24 future climate scenarios. Variables with (N) and(C) indicated models projected within the species native range or at a continental scale, respectively. Results are presented separately for four model sets: the “simple” set of predictors (2 variables in the true niche, 4 predictors per model, 100km buffer), the “complex” set of predictors (3 variables in the true niche, 19 predictors per model, 100km buffer), the “large background” study region (same simulation settings as “complex” but with a 1000km buffer), and the “geographically structured” model set, for which models were constructed and evaluated using geographically partitioned data (same simulations settings as “complex” but with geographic partitioning of data instead of random holdouts).

| Independent variable | Dependent variable | Simple | Complex | Large BG | Geographic |
| --- | --- | --- | --- | --- | --- |
| Test AUC | Spearman (N) | +,.01 |  | +,.02 | +,.08 |
| Test AUC | Spearman (C) |  |  |  | +,.02 |
| Test Max TSS | Spearman (N) | +,.01 |  | +,.02 | +,.07 |
| Test Max TSS | Spearman (C) |  | -,.01 |  | +,.01 |
| Test Max Kappa | Spearman (N) |  | -,.01 | +,.01 | +,.06 |
| Test Max Kappa | Spearman (C) | -, .01 | -,.01 | -,.01 | +,.01 |
| Test AUC | Spearman (F, N) | +,.10 | +,.11 | +,.11 | +,.12 |
| Test AUC | Spearman (F, C) | +,.01 |  | +,.08 | +,.01 |
| Test Max TSS | Spearman (F, N) | +,.09 | +,.10 | +,.11 | +,.12 |
| Test Max TSS | Spearman (F, C) |  |  |  | +,.01 |
| Test Max Kappa | Spearman (F, N) | +,.20 | +,.05 | +,.11 | +,.10 |
| Test Max Kappa | Spearman (F, C) |  |  |  | +,.01 |

**Table 2.**
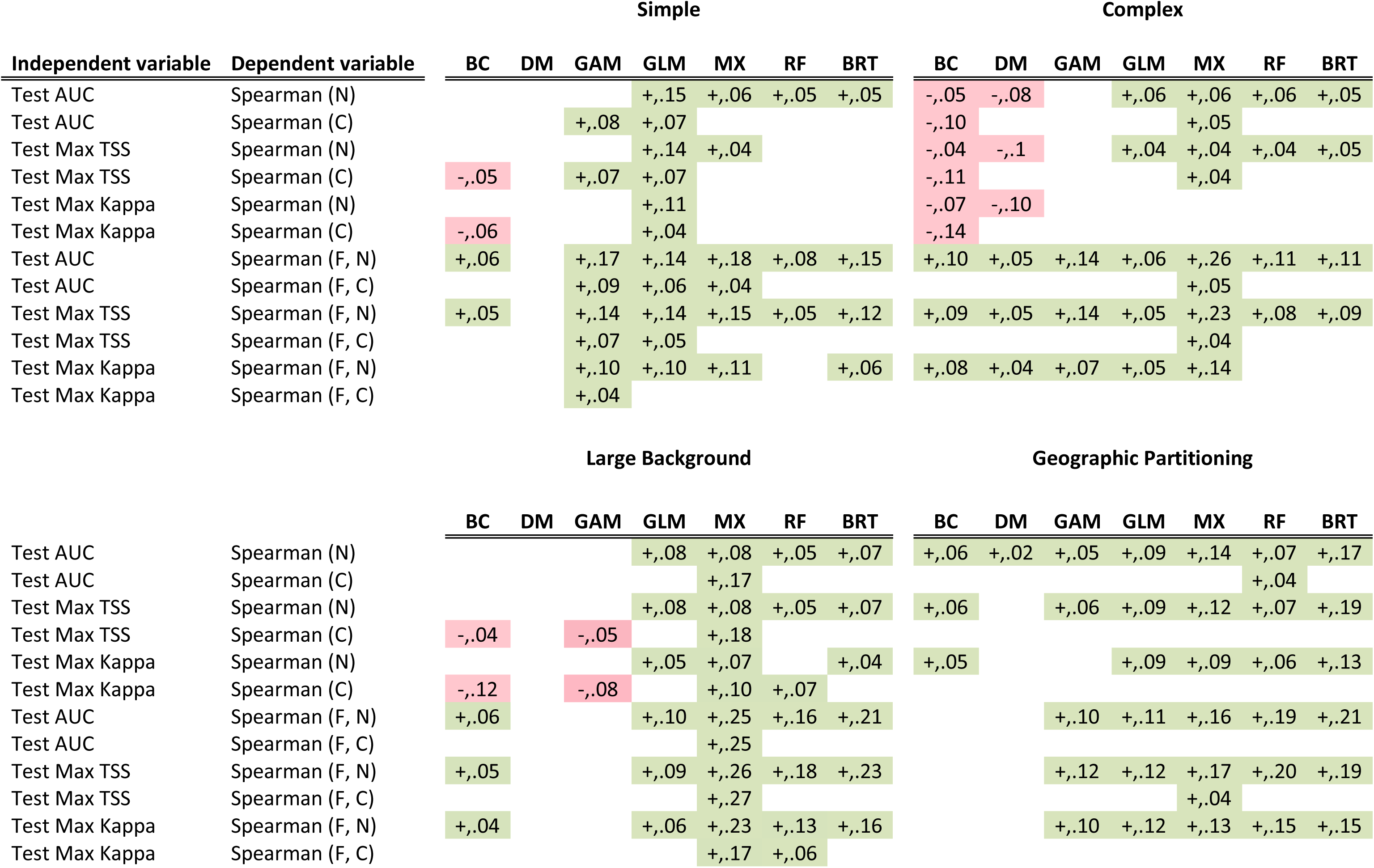
Relationship between discrimination accuracy and functional accuracy, methods considered separately. Significant positive correlations are represented by “+” and green cell color, negative correlations by “-“ and pink cell color. Numbers indicate r^2^ values for each regression. Variables accompanied by (F) indicate that they were measured on models projected across 24 future climate scenarios. Variables with (N) and(C) indicated models projected within the species native range or at a continental scale, respectively. Results are presented separately for four model sets: the “simple” set of predictors (2 variables in the true niche, 4 predictors per model, 100km buffer), the “complex” set of predictors (3 variables in the true niche, 19 predictors per model, 100km buffer), the “large background” study region (same simulation settings as “complex” but with a 1000km buffer), and the “geographically structured” model set, for which models were constructed and evaluated using geographically partitioned data (same simulations settings as “complex” but with geographic partitioning of data instead of random holdouts).

| Independent variable | Dependent variable | Simple |  |  |  |  |  |  | Complex |  |  |  |  |  |  |
| --- | --- | --- | --- | --- | --- | --- | --- | --- | --- | --- | --- | --- | --- | --- | --- |
|  |  | BC | DM | GAM | GLM | MX | RF | BRT | BC | DM | GAM | GLM | MX | RF | BRT |
| Test AUC | Spearman (N) |  |  |  | +,.15 | +,.06 | +,.05 | +,.05 | -,.05 | -,.08 |  | +,.06 | +,.06 | +,.06 | +,.05 |
| Test AUC | Spearman (C) |  |  | +,.08 | +,.07 |  |  |  | -,.10 |  |  |  | +,.05 |  |  |
| Test Max TSS | Spearman (N) |  |  |  | +,.14 | +,.04 |  |  | -,.04 | -,.1 |  | +,.04 | +,.04 | +,.04 | +,.05 |
| Test Max TSS | Spearman (C) | -,.05 |  | +,.07 | +,.07 |  |  |  | -,.11 |  |  |  | +,.04 |  |  |
| Test Max Kappa | Spearman (N) |  |  |  | +,.11 |  |  |  | -,.07 | -,.10 |  |  |  |  |  |
| Test Max Kappa | Spearman (C) | -,.06 |  |  | +,.04 |  |  |  | -,.14 |  |  |  |  |  |  |
| Test AUC | Spearman (F, N) | +,.06 |  | +,.17 | +,.14 | +,.18 | +,.08 | +,.15 | +,.10 | +,.05 | +,.14 | +,.06 | +,.26 | +,.11 | +,.11 |
| Test AUC | Spearman (F, C) |  |  | +,.09 | +,.06 | +,.04 |  |  |  |  |  |  | +,.05 |  |  |
| Test Max TSS | Spearman (F, N) | +,.05 |  | +,.14 | +,.14 | +,.15 | +,.05 | +,.12 | +,.09 | +,.05 | +,.14 | +,.05 | +,.23 | +,.08 | +,.09 |
| Test Max TSS | Spearman (F, C) |  |  | +,.07 | +,.05 |  |  |  |  |  |  |  | +,.04 |  |  |
| Test Max Kappa | Spearman (F, N) |  |  | +,.10 | +,.10 | +,.11 |  | +,.06 | +,.08 | +,.04 | +,.07 | +,.05 | +,.14 |  |  |
| Test Max Kappa | Spearman (F, C) |  |  | +,.04 |  |  |  |  |  |  |  |  |  |  |  |

|  |  | Large Background |  |  |  |  |  |  | Geographic Partitioning |  |  |  |  |  |  |
| --- | --- | --- | --- | --- | --- | --- | --- | --- | --- | --- | --- | --- | --- | --- | --- |
|  |  | BC | DM | GAM | GLM | MX | RF | BRT | BC | DM | GAM | GLM | MX | RF | BRT |
| Test AUC | Spearman (N) |  |  |  | +,.08 | +,.08 | +,.05 | +,.07 | +,.06 | +,.02 | +,.05 | +,.09 | +,.14 | +,.07 | +,.17 |
| Test AUC | Spearman (C) |  |  |  |  | +,.17 |  |  |  |  |  |  |  | +,.04 |  |
| Test Max TSS | Spearman (N) |  |  |  | +,.08 | +,.08 | +,.05 | +,.07 | +,.06 |  | +,.06 | +,.09 | +,.12 | +,.07 | +,.19 |
| Test Max TSS | Spearman (C) | -,.04 |  | -,.05 |  | +,.18 |  |  |  |  |  |  |  |  |  |
| Test Max Kappa | Spearman (N) |  |  |  | +,.05 | +,.07 |  | +,.04 | +,.05 |  |  | +,.09 | +,.09 | +,.06 | +,.13 |
| Test Max Kappa | Spearman (C) | -,.12 |  | -,.08 |  | +,.10 | +,.07 |  |  |  |  |  |  |  |  |
| Test AUC | Spearman (F, N) | +,.06 |  |  | +,.10 | +,.25 | +,.16 | +,.21 |  |  | +,.10 | +,.11 | +,.16 | +,.19 | +,.21 |
| Test AUC | Spearman (F, C) |  |  |  |  | +,.25 |  |  |  |  |  |  |  |  |  |
| Test Max TSS | Spearman (F, N) | +,.05 |  |  | +,.09 | +,.26 | +,.18 | +,.23 |  |  | +,.12 | +,.12 | +,.17 | +,.20 | +,.19 |
| Test Max TSS | Spearman (F, C) |  |  |  |  | +,.27 |  |  |  |  |  |  | +,.04 |  |  |
| Test Max Kappa | Spearman (F, N) | +,.04 |  |  | +,.06 | +,.23 | +,.13 | +,.16 |  |  | +,.10 | +,.12 | +,.13 | +,.15 | +,.15 |
| Test Max Kappa | Spearman (F, C) |  |  |  |  | +,.17 | +,.06 |  |  |  |  |  |  |  |  |

We choose to focus on functional accuracy here instead of calibration for several reasons; first, the application of SDMs more often relies on the relative suitability of habitat than estimating the exact probability of observing a species in a particular place, and functional accuracy more directly estimates this aspect of model behavior. Second, it is already known that discrimination accuracy may be poorly correlated to calibration even when the model gets the relative ranking of habitat right. For example, if the estimated suitability of habitat is a transformation of the true suitability of habitat that preserves relative rankings but not the magnitude of differences in suitability scores (Reineking and Schröder 2006), both discrimination and functional accuracy would be high, but calibration would be poor. Finally, the link between discrimination accuracy and calibration is known to be severely affected by prevalence (Reineking and Schröder 2006, Elith and Graham 2009), but the link between discrimination accuracy and functional accuracy as measured here would not be so affected.

To summarise the relationships between discrimination and functional accuracy for all algorithms considered together (Table 1 and Appendix S3 and S4), we used generalized linear mixed models, and evaluated correlations using McFadden’s pseudo-r^2^ (McFadden 1974). For the remainder of the regressions, we used linear models and the standard coefficient of determination, r^2^. We applied Bonferroni corrections to compensate for problems arising from multiple testing. For these purposes we defined four families of test that we consider independently. Those examining the relationship between discrimination and functional accuracy at each combination of algorithm and complexity level (Tables 1 and 2, n = 12 comparisons per set), and the remainder, which are intended primarily to examine which factors impact overall model quality and as a check to establish that expected relationships between metrics are seen in the simulation results (Appendix S3, n = 11).

Based on results from experiments 1-4, we performed a fifth set of simulation experiments to examine more thoroughly the effects of niche complexity and the number of predictor variables on the relationship between discrimination and functional accuracy (Figure 4). Due to the computational intensity of some SDM algorithms, we restricted analyses to a simpler set of conditions for these simulations. Presence data was generated with no non- target spatial biological processes and no spatial sampling bias, so occurrence points were sampled across the entirety of the suitable habitat. We restricted the modeling process to GLM and Maxent, and only used AUC for evaluating predictions on randomly withheld test data. Simulated niches were built using a number of environmental variables ranging between 1 and 19, and models were inferred with between 1 and 19 variables (2 to 19 for Maxent, due to issues with the software implementation), subject to the constraint that variables that were included within the species’ niche were selected first during model construction. For each combination of number of niche axes and environmental predictors, we performed 300 separate simulations, resulting in 108,300 total simulations per modeling method. For each simulation we recorded the test AUC and the rank correlation between the inferred and true suitability of habitat. For each combination of number of niche axes and predictors, we then measured the rank correlation between discrimination accuracy and functional accuracy across the set of 300 models. This resulted in a metric ranging from 1, where test AUC was a perfect indicator of functional accuracy, to −1, where test AUC was negatively associated with functional accuracy. We fit a linear model to these results which included the number of variables used for the simulated niche, the number of variables used for model construction, and an interaction term.

**Figure 4.**
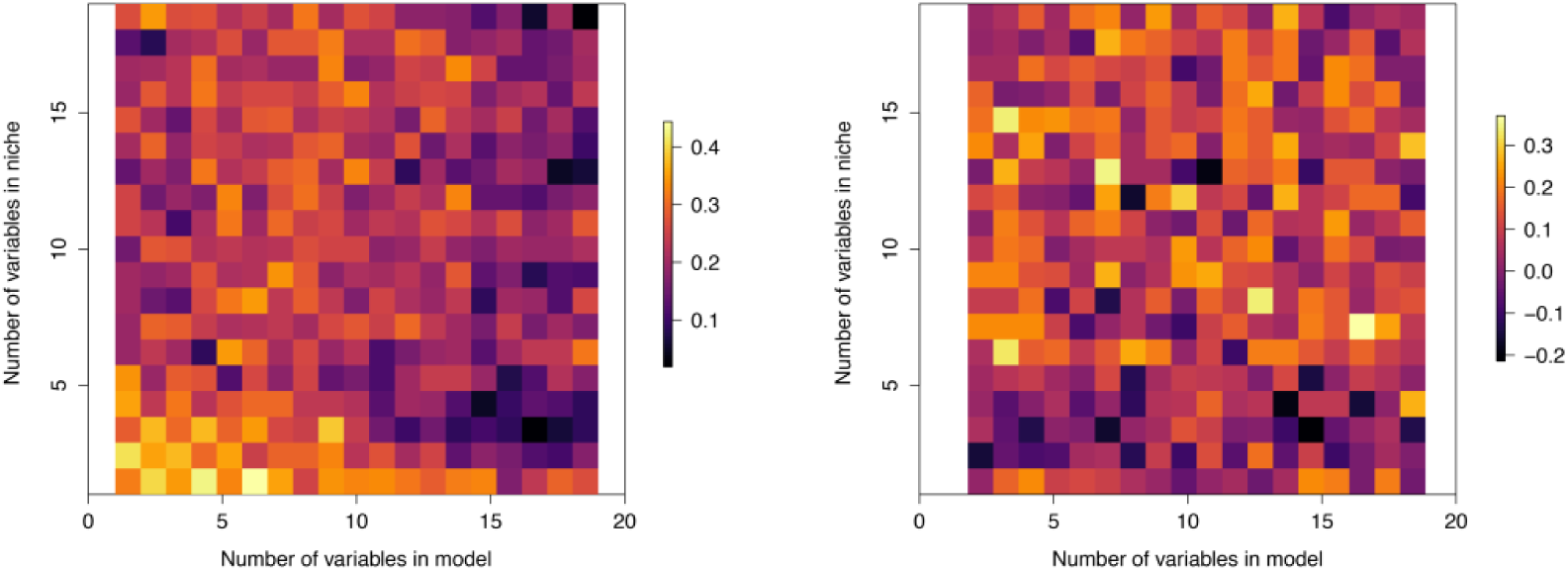
Relationship between number of variables in simulated niche, number of variables in model, and the ability of discrimination to infer functional accuracy for GLM (left) and Maxent (right). Each grid cell represents the output of 300 simulations. The color of each grid cell represents the Spearman rank correlation between test AUC values and functional accuracy.

## Results

Regression outputs for experiments 1-4 are summarized in tables 1 (algorithms pooled) and 2 (algorithms analyzed separately), and also in appendices S3, S4, and S5. TSS, kappa, and AUC were all highly correlated with each other, so we will not discuss them separately. We found that discrimination accuracy on training and test data were correlated, and that functional accuracy in the training region was correlated with functional accuracy outside the training region. This indicates that models that perform well at discrimination accuracy tend to do so regardless of whether it is measured on training or test data, and the same is true of models that perform well at functional accuracy. Functional accuracy was generally fairly good; a majority of models produced estimates of habitat suitability that were positively correlated with the true suitability of habitat, whether measured in the training region (73.8%), or projected to the continental scale (80.8%). However, models performed somewhat worse when projected into future climate scenarios (65.7% were positively correlated with true suitability within the species range, 71.0% at the continental scale).

When all algorithms were analyzed together in a single GLM, discrimination accuracy was a very poor predictor of functional accuracy in all cases (Table 1). Although 31/48 regressions were statistically significant, five were negative correlations, and none had an r^2^ value greater than 0.2. This indicates poor performance of discrimination accuracy at selecting models when comparing between algorithms.

Results of regressions conducted for each algorithm separately are presented in Table2. For all experiments, we find that discrimination accuracy is uninformative or actively misleading about models’ functional accuracy in a majority of cases (significant positive correlations were seen for less than half of comparisons in any simulation experiment). A majority of these correlations were also quite weak; the average r^2^ value was 0.08 (range −.15 to .26). This indicates poor performance of discrimination accuracy at selecting models with high functional accuracy even when comparing models that were built using the same methods. Discrimination accuracy had no negative correlations with functional accuracy when test data were chosen based on a geographic partition of the species’ range, but was still a poor predictor of functional accuracy (average r^2^ = 0.12).

We note an interesting phenomenon here with respect to the size of the buffer regions used to draw background data for model fitting and evaluation; the models built using the largest (1000 km) buffers around occurrence points performed very well, with the highest levels of functional accuracy and discrimination accuracy (Appendix S5). These differences were most prominent for discrimination accuracy, reinforcing previous findings showing that discrimination accuracy is sensitive to study area (Lobo, Jimenez-Valverde et al. 2008). Some previous studies have suggested the models perform best when constructed using fairly small study regions, however those studies have largely assessed model quality via discrimination accuracy within the species’ native range (Acevedo, Jimenez-Valverde et al. 2012, Zhu, Rédei et al. 2014). These results indicate that the relationship between study area size and model performance may be more complex than previously reported, and optimal choices may depend on the applications for which models are designed.

Our fifth experiment examined the effects of complexity across a broader range of complexity values, and found that the ability of AUC to select GLM models with high functional accuracy was negatively correlated with the complexity of the simulated niche and the number of predictor variables. For Maxent models the relationship between discrimination accuracy and functional accuracy (Figure 4) was positively but weakly correlated (r^2^ = .09) with the number of variables in the true niche, but uncorrelated with the number of variables included in the modeling process.

## Discussion

SDM methods are used for many applications in which niche estimates are needed but experimental approaches are impractical. Results for experiments 1-4 demonstrate that many of these models can provide useful estimates of the relative suitability of habitat, the ability of species to invade new areas, and the effects of climate change. However, one of the key steps in any modeling study is the identification of which models from a candidate set perform well and which perform poorly. Our results indicate that the most widely used methods for selecting models are largely uninformative for studies where the goal is to make continuous estimates of habitat suitability, or to estimate the species’ response to an environmental gradient. When algorithms were analyzed separately, 15/149 statistically significant correlations between discrimination and functional accuracy were negative. In these cases discrimination metrics were not just uninformative, but in fact positively misleading for applications where the goal of SDM is to predict the relative suitability of habitat.

In our fifth experiment we examined the effects of niche and model complexity for GLM models across a broader set of conditions, and found that discrimination accuracy predicts functional accuracy only when both the niche and the environmental space it is being modeled in are far simpler than those seen typically in the empirical literature (Figure 4). Even at low levels of complexity, the relationship between discrimination and functional accuracy for GLM is fairly weak (Spearman rank correlation = 0.31 for a single niche axis and predictor variable), and declines rapidly as models become more complex, becoming minimally informative as models approach levels of complexity that are often seen in the empirical literature.

For Maxent models the number of predictors used to model the niche had no effect on the utility of discrimination accuracy for model selection, but there was a weak positive effect of the number of variables used to simulate the true niche (β = .006, p < .05). We hypothesize that the lack of effect of the number of predictors for Maxent is due to its ability to automatically penalize overparameterization; many of the predictors supplied to the algorithm may ultimately have little or no weight in the model. We also note that the most reliable correlations between discrimination accuracy and functional accuracy seen in our simulation results were for Maxent models (Table 2), as would be expected if model complexity and number of predictors were partly responsible for driving the poor performance of discrimination accuracy.

Discrimination accuracy was generally a better predictor of functional accuracy for GLM, GAM, and Maxent models than for the other methods of model construction explored in this study. This is likely due to the internal structure of these models. The simulation approach taken here uses a logistic function to generate sampling probabilities based on a simulated niche, which is composed of smooth (linear or quadratic) responses to a set of environmental variables. As such, the function underlying habitat suitability lies within the set of functions that may be exactly estimated by GLM, GAM, or Maxent, so that estimation of simulated niches is considerably more tractable for those methods. We therefore caution users to refrain from interpreting these results as an endorsement of any particular method when constructing SDMs using empirical data. Rather, we suggest that these results indicate that choice of modeling methods should ideally include intuition or data regarding the potential functional relationship between the environmental predictors and the suitability of habitat. If the functional relationships that may be estimated by an algorithm differ significantly from the true functional relationship, discrimination accuracy is largely uninformative or misleading about models’ ability to predict habitat suitability. This does not necessarily imply that models built using different functional shapes from the true niche are poor estimates of habitat suitability; rather it indicates that discrimination accuracy is uninformative for selecting models with high functional accuracy under these conditions.

Our results clearly indicate that most empirical studies using SDM methods should ideally not rely solely on prediction of withheld occurrence data to assess model quality. However, they also indicate a much more systemic problem for the SDM literature: decades of methodological work in this field have resulted in a set of widely-adopted “best practices”, but a great majority of these studies have focused on optimizing models’ discrimination accuracy on withheld occurrence data from real species distributions (Guisan, Graham et al. 2007, Wisz, Hijmans et al. 2008, Domisch, Kuemmerlen et al. 2013, Boria, Olson et al. 2014, Radosavljevic and Anderson 2014, Moreno-Amat, Mateo et al. 2015, Garcia-Callejas and Araujo 2016, Huang and Frimpong 2016, Kuebler, Hildebrandt et al. 2016, Lopatin, Dolos et al. 2016, Rovzar, Gillespie et al. 2016, Soley-Guardia, Gutierrez et al. 2016). Given the disconnect seen here between discrimination and functional accuracy, it is entirely possible that the “best practices” advocated in these studies have negligible, or even detrimental, effects on model quality for applications where functional accuracy is the goal.

In order to accurately assess the ability of different methods to achieve useful levels of functional accuracy, we argue that the methodological literature must reevaluate its “best practices” via simulations where true habitat suitability and niche parameters are known. While some simulation studies are already being conducted (Meynard, Leroy et al. 2019), these have typically been done in the context of optimizing discrimination accuracy, and as such may also be largely uninformative about estimating habitat suitability as a function of environmental gradients. There are many common practices and assumptions in the field that may need to be reevaluated based on their ability to estimate habitat suitability; choice of algorithm, methods for choosing predictor variables, choice of study area, rarefaction of data, and optimal model complexity are obvious candidates.

In addition, we argue that practitioners must recognize that favoring models based strictly on their spatial predictions is simply inappropriate for many applications. In studies where the goal is to estimate the niche (i.e., maximize functional accuracy), users must become comfortable with the idea that a biologically accurate model may produce relatively poor estimates of species’ current spatial distributions. This is not simply a methodological point brought to light by the current simulation study; it is necessarily true given the existence of non-target phenomena that themselves have spatial structure (e.g., biotic interactions, dispersal). This has been known for years (Jackson and Overpeck 2000, Soberon and Peterson 2005, Anderson 2012, Warren 2012, Warren 2013), yet has been largely ignored in the continued pursuit of methods that produce tighter and tighter fits to training or test data in geographic space.

Investigators familiar with SDM methods will no doubt wish to critically examine the methods used here to infer models; there are other algorithms available, and there are many modeling choices that we did not explore in great depth. However, these criticisms are largely irrelevant to the primary results of this study; while it is certainly possible that greater effort in exploring the space of model choices might improve the accuracy of models, we note that (1) evaluation metrics on randomly withheld test data for the models generated here are not unusual for the range seen in the empirical SDM literature (e.g., Appendix S5), (2) the overall performance of SDM methods is irrelevant to whether or not discrimination accuracy is a valid indicator of functional accuracy, and (3) most SDM users’ methodological preferences are currently chosen based on studies that seek to maximize the very performance metrics that the current study demonstrates are not useful for estimating functional accuracy.

We acknowledge the possibility that there is some subset of modeling approaches not addressed here for which discrimination and functional accuracy are highly correlated. It would be both gratifying and very useful to find such a set of conditions, and that topic deserves to be examined in great depth. However, even if such a set can be found it does not invalidate the conclusion presented here; that there is a large range of modeling algorithms and approaches for which the correlation between discrimination accuracy and functional accuracy is not strong enough to be useful in model selection for many purposes. Similarly, we acknowledge that the disconnect between functional accuracy and discrimination seen here may be affected by sample size, but the sample sizes used here (75 training, 25 test) are not atypical for the ENM literature.

In summary, we demonstrate that, under a broad range of conditions, the ability of a model to successfully predict withheld occurrence data within the training region does not reliably measure its ability to estimate the relationship between environmental gradients and habitat suitability. Discrimination accuracy may be a reasonable metric when the goal is to guide further sampling of occurrences within a species’ current range, without regard for whether the model estimates the true environmental niche or the relative suitability of habitat well. However, this is not often the goal of empirical model construction in the SDM literature.

As a result, the applied and methodological literature in this field are largely based on metrics that may be irrelevant to the intended applications of many models. If the field is to continue to attempt to use SDMs to infer species’ responses to environmental gradients, we must develop methods for model construction and metrics for model evaluation that are more relevant to the actual goals of the modeling process. While we find that geographically structured partitioning of test data does offer some advantages over randomly withheld data, it is clear from this study that even those methods have very limited ability to identify models that accurately estimate the relative suitability of habitat.

We would like to particularly highlight the implications of our results for the development of new methods in this field in the coming years. Many investigators are currently developing methods that incorporate more biological and statistical realism into the SDM process, including the integration of physiological and trait data (Pollock, Kelly et al. 2018) and explicit models of bias (Robinson, Ruiz-Gutierrez et al. 2018), dispersal (Zurell 2017), plasticity (Bush, Mokany et al. 2016), and evolutionary history (Smith, Godsoe et al. 2019). In any system affected by non-target spatial phenomena, these methods will often produce poorer estimates of species’ geographic distributions precisely because they provide better estimates of the environmental niche. We hope that the results presented here will compel the field to evaluate these new methods based on their ability to infer the biological phenomena of interest, as demonstrated using simulations or physiological data, rather than simply reject them due to poor discrimination accuracy on misleading occurrence data.

We feel it is necessary to specifically address one interpretation of these results that we feel is not appropriate: the work presented here is not intended to suggest that any particular method of SDM construction is inherently better or worse than others. While the relative performance of different methods is a very interesting question and one that deserves further exploration within a simulation framework, this study was not designed to address those questions and it would be inappropriate to interpret these results as such. We emphasize that most of the models built from these simulated species were arguably publishable distribution estimates, and were at least somewhat useful as estimates of the species’ niche. Rather, this study is intended to examine the performance of widely-used methods of model selection, and it is those methods that are performing poorly. We demonstrate that we can make both good distribution estimates and good niche estimates using common methods, and in fact produced many models that are good for both purposes. However, our results indicate that we have a difficult time distinguishing good models from bad when our goal is functional accuracy.

At minimum, our results suggest that any empirical study using discrimination accuracy to assess model quality should start with two crucial steps: (1) use a minimal set of predictor variables for which there is an a priori reason to expect that they limit the suitability of habitat for the species, and (2) select algorithms capable of inferring functional responses that are plausible estimates of the underlying biology (e.g., not using a step function in situations where suitability is expected to be a continuous function of the predictor variable). In a sense, these findings are unsurprising; they recapitulate longstanding best practices in the broader literature regarding statistical modeling (Anderson and Burnham 2004, Burnham and Anderson 2004, Gelman and Hill 2006, Zuur, Ieno et al. 2009). However, here we show that failure to make these choices appropriately does not necessarily lead to poor predictions; instead it means that we are largely unable to distinguish good models from bad using species occurrence data. Under these conditions any preference for a given model based on discrimination accuracy may be little better than choosing a model at random.

## Supplementary materials

Appendix S1. Frequency of use of model fit metrics (columns) and partitioning scheme for test data (rows) from a survey of 94 recent applied SDM studies.

Appendix S2. Literature review for metrics of model fit.

Appendix S3. Relationship between evaluation metrics and simulation settings, all algorithms considered together.

Appendix S4. Relationship between discrimination accuracy and functional accuracy using Pearson product moment correlations.

Appendix S5. Discrimination and functional accuracy performance for each simulation experiment.

## Data Availability

Sample code is available on github here: https://github.com/danlwarren/sim-code-Warren-et-al-2019

## Acknowledgements

This work would not have been possible without the financial contributions of the Macquarie University Department of Biology and a DECRA award from the Australian Research Council.

