## Appendix S1 for "Evaluating species distribution models with discrimination accuracy is uninformative for many applications"

|  | **AUC** | **TSS** | **Kappa** | **AICc** | **Other** |
| --- | --- | --- | --- | --- | --- |
| **None** | 1 |  |  | 2 | 2 |
| **Random** | 76 | 20 | 8 | 2 | 13 |
| **Geographic** | 7 |  |  | 2 | 1 |
| **Temporal** | 1 | 1 |  |  | 1 |

Table S1. Frequency of use of model fit metrics (columns) and partitioning scheme for test data (rows) from a survey of 94 recent applied SDM studies.
