## Appendix S2 for "Evaluating species distribution models with discrimination accuracy is uninformative for many applications"

| **Authors** | **Year** | **Title** | **Metrics** | **Subsetting** |
| --- | --- | --- | --- | --- |
| **Aguilar, et al.** | 2015 | Mapping the stray domestic cat (Felis catus) population in New Zealand: Species distribution modelling with a climate change scenario and implications for protected areas | AUC | random |
| **Alamgir, et al.** | 2015 | Modelling spatial distribution of critically endangered Asian elephant and Hoolock gibbon in Bangladesh forest ecosystems under a changing climate | AUC | random |
| **Allen and Lendemer** | 2016 | Climate change impacts on endemic, high-elevation lichens in a biodiversity hotspot | AUC, AICc | geographic |
| **Aryal, et al.** | 2016 | Predicting the distributions of predator (snow leopard) and prey (blue sheep) under climate change in the Himalaya | AICc | random |
| **Ashraf, et al.** | 2016 | Predicting the Potential Distribution of Olea ferruginea in Pakistan incorporating Climate Change by Using Maxent Model | AUC | random |
| **Assis, et al.** | 2016 | Future climate change is predicted to shift long-term persistence zones in the cold-temperate kelp Laminaria hyperborea | TSS | random |
| **Basher and Costello** | 2016 | The past, present and future distribution of a deep-sea shrimp in the Southern Ocean | AUC | random |
| **Beltramino, et al.** | 2015 | Impact of climate change on the distribution of a giant land snail from South America: predicting future trends for setting conservation priorities on native malacofauna | AUC | random |
| **Bonino, et al.** | 2015 | Climate change and lizards: changing species' geographic ranges in Patagonia | AUC | random |
| **Bosso, et al.** | 2016 | Shedding light on the effects of climate change on the potential distribution of Xylella fastidiosa in the Mediterranean basin | AUC, TSS, AUCdiff | random |
| **Brun, et al.** | 2016 | The predictive skill of species distribution models for plankton in a changing climate | AUC, TSS, positive predictive value (PPV) | temporal |
| **Carvalho, et al.** | 2015 | Ecological Niche Modelling Predicts Southward Expansion of Lutzomyia (Nyssomyia) flaviscutellata (Diptera: Psychodidae: Phlebotominae), Vector of Leishmania (Leishmania) amazonensis in South America, under Climate Change | TSS, Cohen's Kappa | random |
| **Chemura, et al.** | 2016 | Bioclimatic modelling of current and projected climatic suitability of coffee (Coffea arabica) production in Zimbabwe | AUC, visual | random |
| **Cruz-Cardenas, et al.** | 2016 | Potential distribution model of Pinaceae species under climate change scenarios in Michoacan | AUC | random |
| **De La Cruz and Ward** | 2016 | Summer-Habitat Suitability Modeling of Myotis sodalis (Indiana Bat) in the Eastern Mountains of West Virginia | AUC, AICc | random |
| **Duan, et al.** | 2016 | The potential effects of climate change on amphibian distribution, range fragmentation and turnover in China | AUC, TSS | random |
| **Friggens and Finch** | 2015 | Implications of Climate Change for Bird Conservation in the Southwestern US under Three Alternative Futures | AUC, Boyce Index | random |
| **Gama, et al.** | 2016 | Predicting global habitat suitability for Corbicula fluminea using species distribution models: The importance of different environmental datasets | AUC, TSS | random |
| **Gedir, et al.** | 2015 | Effects of climate change on long-term population growth of pronghorn in an arid environment | DIC | none |
| **Georgopoulou, et al.** | 2016 | Predicting species richness and distribution ranges of centipedes at the northern edge of Europe | AUC, TSS | random |
| **Hamer, et al.** | 2015 | Shallow environmental gradients put inland species at risk: Insights and implications from predicting future distributions of Eucalyptus species in South Western Australia | AUC, TSS | random |
| **Hanzlicek, et al.** | 2016 | Bayesian Space-Time Patterns and Climatic Determinants of Bovine Anaplasmosis | AUC | random |
| **Hof and Svahlin** | 2016 | The potential effect of climate change on the geographical distribution of insect pest species in the Swedish boreal forest | AUC | geographic |
| **Hoveka, et al.** | 2016 | Effects of climate change on the future distributions of the top five freshwater invasive plants in South Africa | AUC | random |
| **Hu, et al.** | 2015 | Predicting Impacts of Future Climate Change on the Distribution of the Widespread Conifer Platycladus orientalis | AUC | random |
| **Hu, et al.** | 2015 | Projecting Distribution of the Overwintering Population of Sogatella furcifera (Hemiptera: Delphacidae), in Yunnan, China With Analysis on Key Influencing Climatic Factors | AUC | random |
| **Huang, et al.** | 2016 | Limited transferability of stream-fish distribution models among river catchments: reasons and implications | AUC | random and geographic |
| **Ihlow, et al.** | 2016 | Impacts of Climate Change on the Global Invasion Potential of the African Clawed Frog Xenopus laevis | AUC, TSS, Cohen's Kappa | random |
| **Jackson, et al.** | 2015 | Effects of Climate Change on Habitat Availability and Configuration for an Endemic Coastal Alpine Bird | OOB, AUC, TSS, Cohen's Kappa | random |
| **Jueterbock, et al.** | 2016 | The fate of the Arctic seaweed Fucus distichus under climate change: an ecological niche modeling approach | AICc | none |
| **Kafash, et al.** | 2016 | Ensemble distribution modeling of the Mesopotamian spiny-tailed lizard, Saara loricata (Blanford, 1874), in Iran: an insight into the impact of climate change | AUC | random |
| **Kaloveloni, et al.** | 2015 | Winners and losers of climate change for the genus Merodon (Diptera: Syrphidae) across the Balkan Peninsula | AUC, TSS | random |
| **Kamilar, et al.** | 2016 | Anthropogenic and Climatic Effects on the Distribution of Eulemur Species: An Ecological Niche Modeling Approach | AUC | random |
| **Komac, et al.** | 2016 | Modelization of the Current and Future Habitat Suitability of Rhododendron ferrugineum Using Potential Snow Accumulation | AUC, TSS | random |
| **Koncki and Aronson** | 2015 | Invasion Risk in a Warmer World: Modeling Range Expansion and Habitat Preferences of Three Nonnative Aquatic Invasive Plants | AUC | random |
| **Kuebler, et al.** | 2016 | Assessing the importance of topographic variables for the spatial distribution of tree species in a tropical mountain forest | AUC, TSS | random |
| **Kwon, et al.** | 2015 | Predicting potential impacts of climate change on freshwater fish in Korea | Accuracy, AUC, Cohen's Kappa | random |
| **Kwon, et al.** | 2016 | Modelling Vulnerability and Range Shifts in Ant Communities Responding to Future Global Warming in Temperate Forests | AUC | random |
| **Lauzeral, et al.** | 2015 | The iterative ensemble modelling approach increases the accuracy of fish distribution models | AUC, TSS, Kappa, and "the percentage of misclassified sites" | random |
| **Lawrence, et al.** | 2016 | Predicting the potential environmental suitability for Theileria orientalis transmission in New Zealand cattle using maximum entropy niche modelling | AUC, Boyce Index, AICc | geographic |
| **Leach, et al.** | 2016 | Modelling the influence of biotic factors on species distribution patterns | WAIC, DIC | none |
| **Lezama Ochoa, et al.** | 2016 | Present and future potential habitat distribution of Carcharhinus falciformis and Canthidermis maculata by-catch species in the tropical tuna purse-seine fishery under climate change | AUC | random |
| **Li, et al.** | 2016 | Effects of climate change on potential habitats of the cold temperate coniferous forest in Yunnan province, southwestern China | AUC | random |
| **Luo, et al.** | 2015 | Impacts of climate change on distributions and diversity of ungulates on the Tibetan Plateau | AUC | random |
| **Martinez-Freiria, et al.** | 2016 | Contemporary niche contraction affects climate change predictions for elephants and giraffes | TSS | random |
| **Masembe, et al.** | 2016 | Projections of Climate-induced Future Range Shifts among Fruit Fly (Diptera: Tephritidae) Species in Uganda | AUC | random |
| **Mi, et al.** | 2016 | Climate envelope predictions indicate an enlarged suitable wintering distribution for Great Bustards (Otis tarda dybowskii) in China for the 21st century | AUC, TSS | random |
| **Molloy, et al.** | 2016 | Incorporating Field Studies into Species Distribution and Climate Change Modelling: A Case Study of the Koomal Trichosurus vulpecula hypoleucus (Phalangeridae) | AUC | random |
| **Nabout, et al.** | 2016 | The Impact of Global Climate Change on the Geographic Distribution and Sustainable Harvest of Hancornia speciosa Gomes (Apocynaceae) in Brazil | AUC | random |
| **Nazeri, et al.** | 2015 | A geo-statistical approach to model Asiatic cheetah, onager, gazelle and wild sheep shared niche and distribution in Turan biosphere reserve-Iran | AUC | random |
| **Ovalle-Rivera, et al.** | 2015 | Projected Shifts in Coffea arabica Suitability among Major Global Producing Regions Due to Climate Change | AUC, Cohen’s kappa | random |
| **Padalia, et al.** | 2015 | How climate change might influence the potential distribution of weed, bushmint (Hyptis suaveolens)? | AUC | random |
| **Parida, et al.** | 2015 | Climate change expected to drive habitat loss for two key herbivore species in an alpine environment | AUC, TSS | random |
| **Parsa, et al.** | 2015 | Potential geographic distribution of two invasive cassava green mites | AUC | geographic |
| **Penado, et al.** | 2016 | Spatial distribution modelling reveals climatically suitable areas for bumblebees in undersampled parts of the Iberian Peninsula | AUC | geographic |
| **Penman, et al.** | 2015 | Interactive effects of climate change and fire on metapopulation viability of a forest-dependent frog in south-eastern Australia | AUC, expert opinion | random |
| **Perie, et al.** | 2016 | Dominant forest tree species are potentially vulnerable to climate change over large portions of their range even at high latitudes | AUC | random |
| **Petitpierre, et al.** | 2016 | Will climate change increase the risk of plant invasions into mountains? | AUC, Boyce Index | random |
| **Porfirio, et, al.** | 2016 | Projected direct and indirect effects of climate change on the Swift Parrot, an endangered migratory species | AUC | random |
| **Prieto-Torres, et al.** | 2016 | Response of the endangered tropical dry forests to climate change and the role of Mexican Protected Areas for their conservation | AUC | random |
| **Priti, et al.** | 2016 | Modeling impacts of future climate on the distribution of Myristicaceae species in the Western Ghats, India | AUC | random |
| **Raghavan, et al.** | 2016 | Maximum Entropy-Based Ecological Niche Model and Bio-Climatic Determinants of Lone Star Tick (Amblyomma americanum) Niche | AUC | random |
| **Rahmati, et al.** | 2016 | Application of GIS-based data driven random forest and maximum entropy models for groundwater potential mapping: A case study at Mehran Region, Iran | AUC | random |
| **Rasquinha and Sankaran** | 2016 | Modelling biome shifts in the Indian subcontinent under scenarios of future climate change | Kappa, Accuracy | random |
| **Regos, et al.** | 2016 | Predicting the future effectiveness of protected areas for bird conservation in Mediterranean ecosystems under climate change and novel fire regime scenarios | AUC | random |
| **Rovzar, et al.** | 2016 | Landscape to site variations in species distribution models for endangered plants | AUC | random |
| **Saenz-Romero, et al.** | 2015 | Pinus leiophylla suitable habitat for 1961-1990 and future climate | OOB | random |
| **Sarma, et al.** | 2015 | Effect of Climate Change on Invasion Risk of Giant African Snail (Achatina fulica Ferussac, 1821: Achatinidae) in India | AUC | random |
| **Scales, et al.** | 2016 | Identifying predictable foraging habitats for a wide-ranging marine predator using ensemble ecological niche models | AUC, TSS, Boyce Index | random |
| **Schwalm, et al.** | 2016 | Habitat availability and gene flow influence diverging local population trajectories under scenarios of climate change: a place-based approach | AUC, OOB | random |
| **Sidder, et al.** | 2016 | Using spatiotemporal correlative niche models for evaluating the effects of climate change on mountain pine beetle | AUC | random |
| **Silva, et al.** | 2015 | Range increase of a Neotropical orchid bee under future scenarios of climate change | AUC | random |
| **Silva, et al.** | 2016 | Distributional modeling of Mantophasmatodea (Insecta: Notoptera): a preliminary application and the need for future sampling | AUC, TSS | random |
| **Soley-Guardia, et al.** | 2016 | Are we overestimating the niche? Removing marginal localities helps ecological niche models detect environmental barriers | AUC | geographic |
| **Struebig, et al.** | 2015 | Anticipated climate and land-cover changes reveal refuge areas for Borneo's orangutans | AUC | random |
| **Su, et al.** | 2015 | Climate Change-Induced Range Expansion of a Subterranean Rodent: Implications for Rangeland Management in Qinghai-Tibetan Plateau | AUC | random |
| **Swab, et al.** | 2015 | The role of demography, intra-species variation, and species distribution models in species' projections under climate change | AUC | random |
| **Szczepanska, et al.** | 2016 | Lichen-forming fungi of the genus Montanelia in Poland and their potential distribution in Central Europe | AUC | random |
| **Tainio, et al.** | 2016 | Conservation of grassland butterflies in Finland under a changing climate | AUC, TSS | random |
| **Tarkesh and Jetschke** | 2016 | Investigation of current and future potential distribution of Astragalus gossypinus in Central Iran using species distribution modelling | AUC, Kappa | random |
| **Torres, et al.** | 2015 | Poor Transferability of Species Distribution Models for a Pelagic Predator, the Grey Petrel, Indicates Contrasting Habitat Preferences across Ocean Basins | AUC, percent deviance explained | random |
| **Vasconcelos and Do Nascimento** | 2016 | Potential Climate-Driven Impacts on the Distribution of Generalist Treefrogs in South America | AUC | random |
| **Vaz, et al.** | 2016 | Using ecological niche models to predict the impact of global climate change on the geographical distribution and productivity of Euterpe oleracea Mart. (Arecaceae) in the Amazon | AUC | random |
| **Vergara, et al.** | 2016 | Shaken but not stirred: multiscale habitat suitability modeling of sympatric marten species (Martes martes and Martes foina) in the northern Iberian Peninsula | AUC | random |
| **Weinert, et al.** | 2016 | Modelling climate change effects on benthos: Distributional shifts in the North Sea from 2001 to 2099 | AUC | random |
| **West, et al.** | 2015 | Using High-Resolution Future Climate Scenarios to ForecastBromus tectorum Invasion in Rocky Mountain National Park | AUC | random |
| **Wittmann, et al.** | 2016 | Confronting species distribution model predictions with species functional traits | AUC | random |
| **Wright, et al.** | 2016 | Advances in climate models from CMIP3 to CMIP5 do not change predictions of future habitat suitability for California reptiles and amphibians | AICc | none |
| **Xavier, et al.** | 2016 | Biogeography of Cephalopods in the Southern Ocean Using Habitat Suitability Prediction Models | AUC | random |
| **Xie, et al.** | 2016 | Projecting the Global Distribution of the Emerging Amphibian Fungal Pathogen, Batrachochytrium dendrobatidis, Based on IPCC Climate Futures | AUC | random |
| **Yi, et al.** | 2016 | Maxent modeling for predicting the potential distribution of endangered medicinal plant (H. riparia Lour) in Yunnan, China | AUC | random |
| **Yuan, et al.** | 2015 | Maxent modeling for predicting the potential distribution of Sanghuang, an important group of medicinal fungi in China | AUC | random |
| **Zaidi, et al.** | 2016 | Distribution Modeling of three screwworm species in the ecologically diverse landscape of North West Pakistan | AUC, TSS | random |
| **Zhu, et al.** | 2016 | Mapping the ecological dimensions and potential distributions of endangered relic shrubs in western Ordos biodiversity center | AUC | none |

Table S2. Use of model fit metrics and test data partitioning in the empirical literature, full literature search results.
