## Appendix S3 for "Evaluating species distribution models with discrimination accuracy is uninformative for many applications"

| **Independent variable** | **Dependent variable** | **Simple** | **Complex** | **Large BG** | **Geographic** |
| --- | --- | --- | --- | --- | --- |
| Train AUC | Test AUC | +,.38 | +, .38 | +,.69 | +,.02 |
| Spearman (N) | Spearman (C) | +,.31 | +, .11 | +,.13 | +,.15 |
| Test AUC | Test Max Kappa | +,.70 | +,.70 | +,.77 | +,.69 |
| Test AUC | Test Max TSS | +,.93 | +,.93 | +,.92 | +,.90 |
| Test Max Kappa | Test Max TSS | +,.73 | +,.71 | +,.80 | +,.66 |
| Bias Strength | Spearman (N) |  |  |  |  |
| Prop. Suitable Occupied | Spearman (N) |  |  |  |  |
| Bias Strength | Spearman (C) |  |  | -,.01 |  |
| Prop. Suitable Occupied | Spearman (C) |  |  | +,.02 |  |
| Size of Range | Spearman (N) | -,.14 | -,.10 | -,.11 | -,.10 |
| Size of Range | Spearman (C) | -,.11 | -,.01 | -,.03 | -,.01 |

Table S3.1 Relationship between evaluation metrics and simulation settings, all algorithms considered together. Significant positive correlations are represented by “+” and green cell color, negative correlations by “-“ and pink cell color. Numbers indicate r^2^ values for each regression. Variables accompanied by (F) indicate that they were measured on models projected across 24 future climate scenarios. Variables with (N) and(C) indicated models projected within the species native range or at a continental scale, respectively.
