## Appendix S4 for "Evaluating species distribution models with discrimination accuracy is uninformative for many applications"

| **Independent variable** | **Dependent variable** | **Simple** | **Complex** | **Large BG** | **Geographic** |
| --- | --- | --- | --- | --- | --- |
| Test AUC | Pearson (N) | +,.01 |  | +,.02 | +,.11 |
| Test AUC | Pearson (C) | +,.03 |  | +,.04 | +,.11 |
| Test Max TSS | Pearson (N) |  |  | +,.02 | +,.09 |
| Test Max TSS | Pearson (C) | +,.02 |  | +,.06 | +,.09 |
| Test Max Kappa | Pearson (N) |  | -,.01 | +,.01 | +,.08 |
| Test Max Kappa | Pearson (C) |  | -,.01 | +,.02 | +,.07 |

Table S4.1 Relationship between discrimination accuracy and functional accuracy, using Pearson product moment correlation to measure functional accuracy, with all modeling algorithms pooled. Significant positive correlations are represented by “+” and green cell color, negative correlations by “-“ and pink cell color. Numbers indicate r^2^ values for each regression.

|  |  |  | **Simple** | | | | | | |  | **Complex** | | | | | | |
| --- | --- | --- | --- | --- | --- | --- | --- | --- | --- | --- | --- | --- | --- | --- | --- | --- | --- |
| **Independent variable** | **Dependent variable** |  | **BC** | **DM** | **GAM** | **GLM** | **MX** | **RF** | **BRT** |  | **BC** | **DM** | **GAM** | **GLM** | **MX** | **RF** | **BRT** |
| Test AUC | Pearson (N) |  | -,.07 | -,.12 |  | +,.18 | +,.04 | +,.06 | +,.05 |  | -,.06 | -,.12 |  | +,.13 | +,.04 | +,.07 | +,.04 |
| Test AUC | Pearson (C) |  |  |  | +,.10 | +,.16 | +,.11 | +,.18 | +,.13 |  |  |  | +,.04 |  | +,.09 | +,.11 | +,.11 |
| Test Max TSS | Pearson (N) |  | -,.07 | -,.13 |  | +,.18 |  |  |  |  | -,.05 | -,.15 |  | +,.11 |  | +,.05 | +,.05 |
| Test Max TSS | Pearson (C) |  |  |  | +,.09 | +,.14 | +,.10 | +,.12 | +,.14 |  |  |  | +,.04 |  | +,.07 | +,.10 | +,.10 |
| Test Max Kappa | Pearson (N) |  | -,.05 | -,.12 |  | +,.12 |  |  |  |  | -,.09 | -,.14 |  | +,.07 |  |  |  |
| Test Max Kappa | Pearson (C) |  |  |  |  |  |  |  |  |  |  |  |  |  |  |  |  |
|  |  |  | **Large Background** | | | | | | |  | **Geographic Partitioning** | | | | | | |
|  |  |  | **BC** | **DM** | **GAM** | **GLM** | **MX** | **RF** | **BRT** |  | **BC** | **DM** | **GAM** | **GLM** | **MX** | **RF** | **BRT** |
| Test AUC | Pearson (N) |  |  | -,.07 | +,.06 | +,.17 | +,.06 | +,.06 | +,.13 |  | +,.07 |  | +,.13 | +,.13 | +,.17 | +,.07 | +,.23 |
| Test AUC | Pearson (C) |  |  |  | +,.08 |  | +,.12 | +,.12 | +,.18 |  | +,.11 | +,.05 | +,.04 | +,.06 | +,.18 | +,.19 | +,.23 |
| Test Max TSS | Pearson (N) |  |  | -,.06 | +,.05 | +,.17 | +,.06 | +,.07 | +,.13 |  | +,.07 |  | +,.13 | +,.13 | +,.14 | +,.06 | +,.22 |
| Test Max TSS | Pearson (C) |  |  |  | +,.08 |  | +,.10 | +,.12 | +,.17 |  | +,.12 | +,.05 | +,.05 | +,.05 | +,.17 | +,.21 | +,.26 |
| Test Max Kappa | Pearson (N) |  |  | -,.09 |  | +,.08 |  |  | +,.06 |  | +,.04 |  | +,.09 | +,.12 | +,.11 | +,.06 | +,.20 |
| Test Max Kappa | Pearson (C) |  |  |  |  |  |  |  | +,.07 |  | +,.08 |  | +,.03 | +,.05 | +,.09 | +,.12 | +,.14 |

Table S4.2 Relationship between discrimination accuracy and functional accuracy, using Pearson product moment correlation to measure functional accuracy, with modeling algorithms considered separately. Significant positive correlations are represented by “+” and green cell color, negative correlations by “-“ and pink cell color. Numbers indicate r^2^ values for each regression.
