## Appendix S5 for "Evaluating species distribution models with discrimination accuracy is uninformative for many applications"

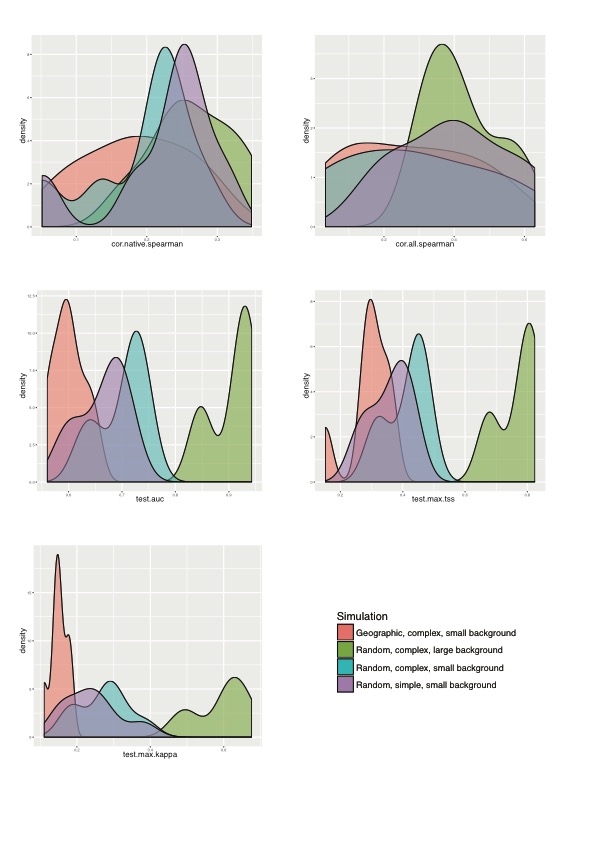

Figure S5.1 Average performance of models. A majority of models made predictions positively correlated with the suitability of habitat, both in the native range (top left) and when projected to the continental scale (top right). Discrimination accuracy for most models was within the range typically seen in the empirical literature. We found that models have considerably higher discrimination accuracy when the geographic background is large, and much lower discrimination accuracy when discrimination accuracy is tested on spatially partitioned data. Both of these results are consistent with existing empirical studies.
